# Pairwise common variant meta-analyses of schizophrenia with other psychiatric disorders reveals shared and distinct gene and gene-set associations

**DOI:** 10.1101/725614

**Authors:** William R. Reay, Murray J. Cairns

**Author notes:** To whom correspondence should be addressed: Professor Murray Cairns, Medical Sciences Building, University Drive, Callaghan, NSW 2308, Australia.

## Abstract

The complex aetiology of schizophrenia is postulated to share factors with other psychiatric disorders. Recently, this has been supported by genome-wide association studies, with several psychiatric phenotypes displaying high genomic correlation with schizophrenia. We sought to investigate pleiotropy amongst the common variant genomics of schizophrenia and seven other psychiatric disorders using a multimarker test of association. Gene-based analysis of common variation revealed over 50 schizophrenia-associated genes shared with other psychiatric phenotypes; including bipolar disorder, major depressive disorder, ADHD, and autism spectrum disorder. In addition, we uncovered 78 genes significantly enriched with common variant associations for schizophrenia that were not linked to any of these seven disorders (*P* > 0.05). Transcriptomic imputation was then leveraged to investigate the functional significance of variation mapped to these genes, prioritising several interesting functional candidates. Pairwise meta-analysis of schizophrenia and each psychiatric phenotype further revealed 330 significantly associated genes (*P*_Meta_ < 2.7 × 10^−6^) that were only nominally associated with each disorder individually (*P* < 0.05). Multivariable gene-set association suggested that common variation enrichment within biologically constrained genes observed for schizophrenia also occurs across several psychiatric phenotypes. These analyses consolidate the overlap between the genomic architecture of schizophrenia and other psychiatric disorders and uncovered several pleiotropic genes which warrant further investigation.

**AUTHOR SUMMARY:** Schizophrenia and other psychiatric disorders have many similarities, and this includes features of their overall genetic risk. Here, we investigate genes which may play a role in schizophrenia as well one or more of seven other psychiatric phenotypes and demonstrate that a number of them are pleiotropic and influence at least one other disorder. We also identify genes amongst the psychiatric disorders studied here which only show association with schizophrenia. Furthermore, we find a number of genes which were only significant when combining genetic association data from schizophrenia and one of the other seven disorders, suggesting there are shared genetic influences that are revealed through the power of joint analysis. This study identifies interesting novel shared (pleiotropic) genes in psychiatry which warrant future study.

## INTRODUCTION

Schizophrenia is a psychiatric disorder which is proposed to arise from a complex interplay between heritable and environmental factors. As a polygenic disorder, the genomic architecture of schizophrenia encompasses variation throughout the genome with population frequencies ranging from common to ultra-rare (1-4). Genome-wide association studies (GWAS) of schizophrenia have been able to recapitulate a notable proportion of the heritability estimated from twin-studies, with SNP based heritability in the most recent GWAS calculated to be approximately 23% on the liability scale, assuming a population prevalence of 0.7% (1). This polygenic signal is distributed amongst many genes and biological systems genome-wide, and thus, further research is required to fully appreciate the underlying biology captured by GWAS and its relevance to the pathophysiology of schizophrenia.

The diagnostic boundaries between schizophrenia and other psychiatric disorders remain difficult to define. Psychiatric co-morbidities are common in patients with schizophrenia (5), whilst the defined clinical presentation of the disorder itself resembles that of other diagnoses. For instance, major depressive disorder (MDD) diagnosis is prevalent amongst individuals with schizophrenia (6). However, negative symptoms inherent to schizophrenia, which include avolition and asociality, are also closely linked to MDD despite their classification as distinct clinical entities (7). Collectively, this supports the existence of shared aetiological factors between schizophrenia and the spectrum of psychiatric illness.

Genomic evidence reinforces this trans-diagnostic paradigm, facilitated by the GWAS now available for a number of psychiatric disorders in addition to schizophrenia (8-14). Linkage disequilibrium (LD) score regression has demonstrated that schizophrenia displays positive genomic correlation with several psychiatric phenotypes (15), with the most statistically significant relationship observed with bipolar disorder (BIP). The biological mechanisms encapsulated by these cross-disorder correlations remain largely uncharacterised. Genes and systems associated with schizophrenia which exhibit pleiotropy within psychiatry, that is, a significant relationship with another disorder, may be particularly biologically salient. Further, elucidation of genetic factors specific to schizophrenia may aid in identifying the underlying origin of clinical features which are more distinct to the disorder. Previous cross-disorder association analyses have largely focused on individual SNPs and consider a set of psychiatric phenotypes simultaneously (16). In this study, we implemented a multimarker test of association from a schizophrenia focus in relation to seven other psychiatric disorders – bipolar disorder (BIP), attention deficit/hyperactivity disorder (ADHD), autism spectrum disorder (ASD), major depressive disorder (MDD), obsessive compulsive disorder (OCD), Tourette’s syndrome (TS), and eating disorder (ED). We sought to identify pleiotropic genes and gene-sets which were associated with schizophrenia and at least one other phenotype. We uncovered several interesting schizophrenia associated genes and gene-sets which are shared, along with a subset of genes specifically linked to schizophrenia.

## RESULTS

### Gene-based association revealed genes specific to schizophrenia and shared with other psychiatric disorders

We implemented a multimarker method to find genes enriched with common variant associations for each disorder. Considering genes outside the extended major histocompatibility complex (MHC) region, 456 genes were associated with schizophrenia below the Bonferroni threshold (*P* < 2.7 × 10^−6^, Supplementary Table 1). The most significantly associated gene with schizophrenia was *CACNA1C* (*P* = 1.76 × 10^−20^), which encodes the calcium-voltage gated channel subunit alpha 1C. Thereafter, the next three most significant schizophrenia genic-associations were: Dihydropyrimidine dehydrogenase (*DPYD, P* = 1.52 × 10^−19^), Protein phosphatase 1 regulatory subunit 16B (*PPP1R16B, P* = 5.85 × 10^−19^), and Transcription factor 4 (*TCF4, P* = 5.68 × 10^−18^). The seven other psychiatric disorders also all had at least one significant genic-association (Supplementary Table 2), ranging from 121 Bonferroni significant associations for BIP, to just one for OCD (Kit Proto-Oncogene, Receptor Tyrosine Kinase [*KIT*], *P* = 2.3 × 10^−7^).

**Table 1.**
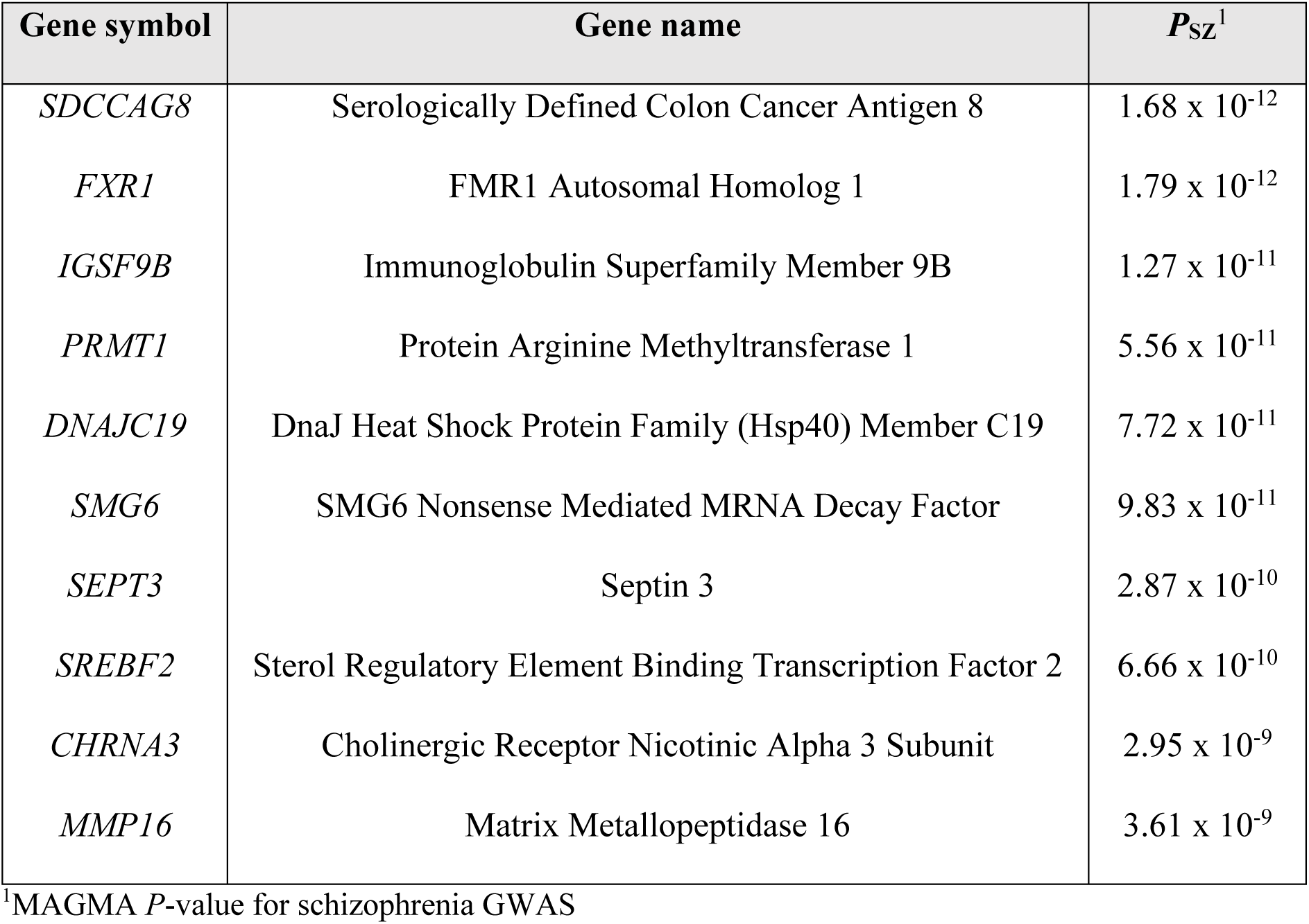
The top ten most significantly associated schizophrenia genes by MAGMA which were not nominally associated (*P* > 0.05) with any of the seven other psychiatric disorders investigated.

We investigated the association of the 456 genes which survived Bonferroni correction in the schizophrenia GWAS in relation to the other seven disorders to identify pleiotropic genes (Fig. 1). In total, there were 67 genes significant after Bonferroni correction in the schizophrenia GWAS and at least one additional psychiatric GWAS. *CACNA1C* was the most significant pleiotropic gene in terms of association with schizophrenia, as it also survived correction for the BIP GWAS, *P*_BIP_ = 1.72 × 10^−9^). BIP shared the greatest number of significant genes with schizophrenia – N_Shared_ = 47, 39% of all significant BIP genes. Significant genes after multiple testing correction were also shared between schizophrenia with ADHD (N_Shared_ = 8, 32% of all significant ADHD genes), MDD (N_Shared_ = 10, 53% of all significant MDD genes), and ASD (N_Shared_ = 3, 18% of all significant ASD genes). One gene, Sortilin related VPS10 domain containing receptor 3 (*SORCS3*), was associated with schizophrenia and two of the additional disorders – ADHD and MDD (*P*_SZ_ = 2.91 × 10^−8^, *P*_ADHD_ = 1.51 × 10^−9^, and *P*_MDD_ = 5.66 × 10^−8^).

**Figure 1.**
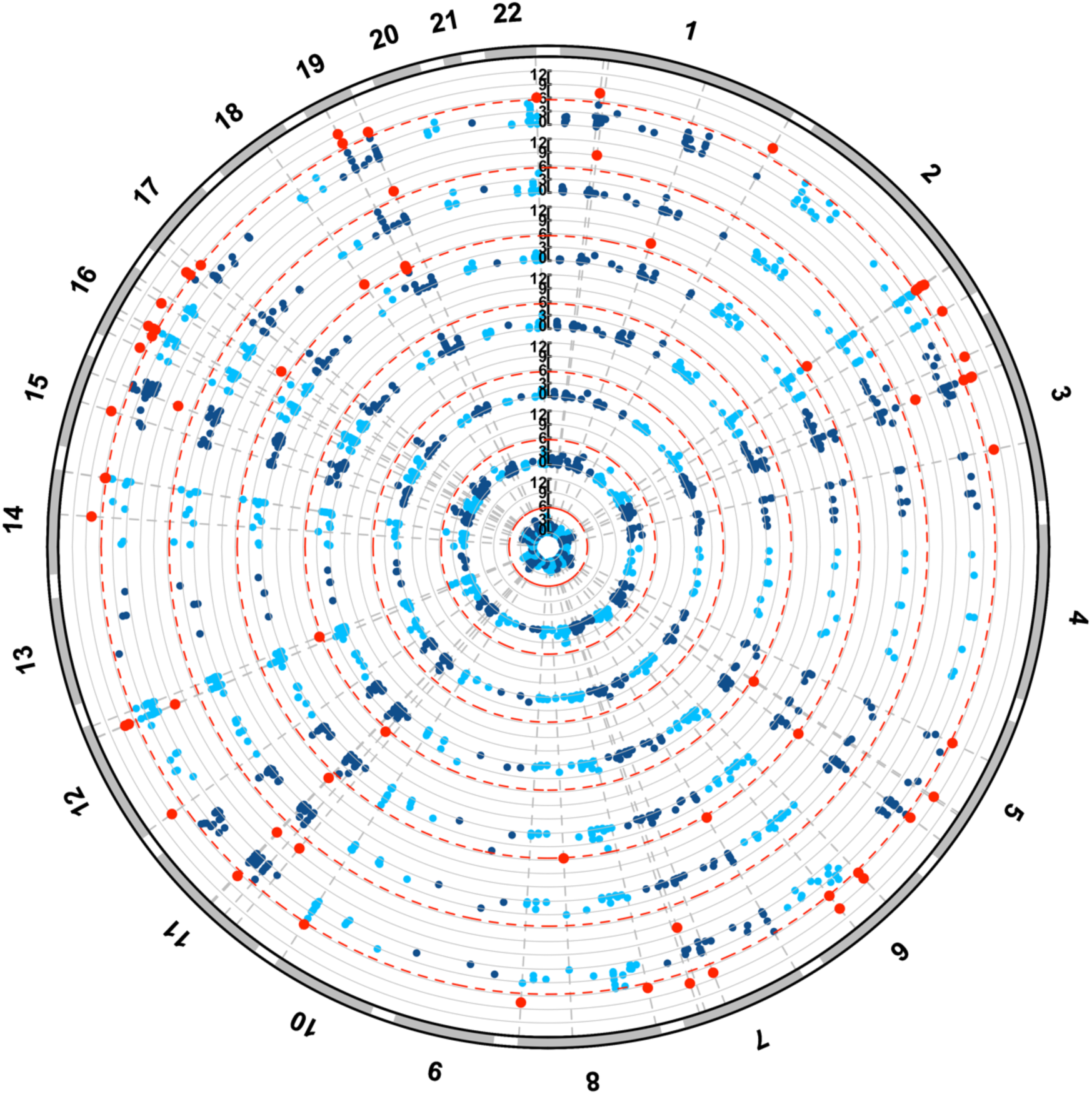
Schizophrenia associated genes shared with other psychiatric disorders. Association of Bonferroni significant schizophrenia genes with 7 other psychiatric disorders. Results presented as a circular Manhattan plot for the −log_10_ *P*-value of association for each gene per disorder. Significant schizophrenia genes by MAGMA which survive correction in each disorder are highlighted red (*P* < 2.7 × 10^−6^). Each shell of the plot represents a different disorder, radiating outwards in the following order: OCD, TS, ED, ASD, MDD, ADHD, and BIP.

A transcriptome wide association study (TWAS) was performed for each of the disorders which displayed at least one shared genic association with schizophrenia (Supplementary Table 3, 4). TWAS leverages imputed models of genetically regulated expression to identify genes for which expression is genetically correlated with the trait of interest (17). We implemented this approach to interpret the direction of effect of these pleiotropic genes in relation to their expression. Specifically, we aimed to identify genes whose predicted expression is associated with schizophrenia and at least one other psychiatric phenotype. Weights were derived from the dorsolateral prefrontal cortex (DLPFC) and blood as described elsewhere (18).

Firstly, we considered models for significant *cis*-heritable genes in the DLPFC; 24 of the 67 candidate pleiotropic genes had available models in this tissue. The imputed expression of three genes was associated with both schizophrenia and BIP (*P* < 9.2 × 10^−6^, corrected for the 5420 models tested). In two of these instances, overexpression was associated with both schizophrenia and BIP: Hyaluronan and proteoglycan link protein 4 [*HAPLN4* – (schizophrenia: *Z*_*TWAS*_ = 5.03, *P* = 4.66 × 10^−7^), (BIP: *Z*_*TWAS*_ = 5.6506, *P* = 1.6 × 10^−8^)], and NIMA related kinase 4 [*NEK4* – (schizophrenia: *Z*_*TWAS*_ = 5.34, *P* = 9.25 × 10^−8^), (BIP: *Z*_*TWAS*_ = 5.13, *P* = 2.9 × 10^−7^)]. An additional gene, *GLT8D1*, proximally located to *NEK4*, also survived correction the schizophrenia and BIP analyses. Conditional analysis was performed to reveal whether these were conditionally independent. *NEK4* was conditionally significant in both disorders below the Bonferroni threshold, whilst in this construct *GLT8D1* was only marginally significant in schizophrenia (*P*_Conditional_ = 0.034) and not significant in BIP (*P*_Conditional_ = 0.69). In addition, conditional analyses for schizophrenia of the *HAPLN4* locus suggested that this association was primarily explained by another gene, *GATAD2A*. Interestingly, *GATAD2A* did not survive correction in the BIP TWAS (*P* = 3.8 × 10^−3^), and implicated *HAPLN4*, in contrast to schizophrenia. There are two plausible explanations for this phenomenon: firstly, technical variability may affect the number of loci tagged in this region between the respective GWAS, or secondly, different biological effects may be conferred by variation mapped to this locus in BIP relative to schizophrenia. Further biological investigation is required to elucidate the mechanisms underlying this signal. An additional four shared genes derived by MAGMA were significant after correction in relation to schizophrenia but trended towards significance for BIP (*P*_BIP_ < 9.25 × 10^−4^: DDHD2, *TMEM110, ITIH4, COG8*), whilst downregulation of *SFMBT1* expression trended towards significance for association with risk for both disorders. Moreover, decreased expression of Mediator complex subunit 8 (*MED8*) was associated with both schizophrenia and ADHD. Imputed expression models derived from blood data were applied as above, with 22 significant models available. Upregulation of Neuromedin B (*NMB*), for which expression was not significantly *cis*-heritable in the brain, was associated with schizophrenia and BIP risk. *GLN3* (G protein nuclear 3) was similarly significant for schizophrenia and BIP, however, due to its proximity to *NEK4* this may arise from the same underlying genomic signal. In blood, *NEK4* does not display adequate *cis*-heritability for imputation, and thus, conditional analyses cannot be directly performed. A pleiotropic effect for overexpression of Zinc Finger DHHC-Type Containing 5 (*ZDHHC5*) on schizophrenia and MDD was also observed using blood SNP weights – schizophrenia: *Z*_*TWAS*_ = 5.96, *P* = 2.49 × 10^−9^, MDD: *Z*_*TWAS*_ = 4.54, *P* = 5.62 × 10^−6^.

### Common variant derived genic-association specific to schizophrenia in psychiatry

We then sought to identify genes which were only associated with schizophrenia (Supplementary Table 5). Firstly, the majority of genes associated with schizophrenia (N = 390, 86% of all genes associated with schizophrenia) did not survive Bonferroni correction for any of the seven other disorders. However, as many of these genes trended towards corrected significance in one or more of the phenotypes, we wanted to identify the subset of genes which were not nominally significant for any other psychiatric GWAS examined in this study (*P* > 0.05, N = 78, 17% of all genes associated with schizophrenia). The gene encoding Serologically Defined Colon Cancer Antigen 8 (*SDCCAG8, P*_SZ_ = 1.68 × 10^−12^) was the most significantly associated gene which did not display nominal association with any of the other psychiatric traits. The top ten of these genes are presented in Table 1. We investigated biological systems for which this subset was overrepresented relative to the other schizophrenia associated genes. No pathways survived multiple testing correction, however, three trended towards corrected significance: *Reactome: Extracellular matrix organisation* (*P* = 2.31 × 10^−3^, *q* = 0.0832), *GO: Cellular response to stimulus* (*P* = 6.82 × 10^−3^, *q* = 0.305), and *GO: Leukocyte proliferation* (*P* = 8.76 × 10^−3^, *q* = 0.305).

TWAS was also performed for this subset of genes using both the brain and blood derived tissue panels. Of the genes tested, 26 and 18 models had sufficient overlapping SNP weights for the analysis considering brain and blood respectively (Supplementary Table 6). Nine genes survived correction across both tissues. The most significant association with schizophrenia in the DLPFC construct was for this subset of genes was increased predicted expression of *SDCCAG8* – *Z*_TWAS_ = 6.16, *P* = 7.21 × 10^−10^. Several of these genes remained significant after the application of both Bonferroni correction and conditional analysis for the DLPFC models – *CLNC3* (*Z*_TWAS_ = 5.36, *P* = 8.36 × 10^−8^), *SLC45A1* (*Z*_TWAS_ = −5.2, *P* = 1.97 × 10^−7^), *KCNN3* (*Z*_TWAS_ = −5.14, *P* = 2.63 × 10^−7^), *GIGYF1* (*Z*_TWAS_ = 4.59, *P* = 4.32 × 10^−6^), and *FAM114A2* (*Z*_TWAS_ = −4.51, *P* = 6.54 × 10^−6^). However, after conditional analysis considering proximally located associations, *SDCCAG8* was only marginally significant (*P* = 0.016), with another gene at this locus (*CEP170*) explaining the majority of the association. In addition, the two genes which survived correction for blood were also only marginally significant after a conditional test (*HVCN1*: *P*_Raw_ = 3.74 × 10^−7^, *P*_Conditonal_ *=* 0.011; *SBNO1*: *P*_Raw_ = 7.78 × 10^−7^, *P*_Conditonal_ *=* 0.025).

### Biologically constrained genes were enriched with common variation across multiple psychiatric disorders

We performed two gene-set association analyses, firstly, two hypothesis driven gene-sets associated with schizophrenia in the largest GWAS after multiple-testing correction were investigated in relation to the psychiatric disorders, and secondly, a data-driven approach using 7296 gene-sets from the MSigDB (19). The hypothesis driven constructs were genes intolerant to loss of function variation (probability of loss of function intolerant [pLI] ≥ 0.9, N_Genes_ = 3230), and targets of the mRNA binding protein fragile X mental retardation protein (FMRP) (1, 20, 21). All disorders displayed some level of enrichment for common variant associations in mutation intolerant gene-set, except for ED and OCD (Figure 2).

**Figure 2.**
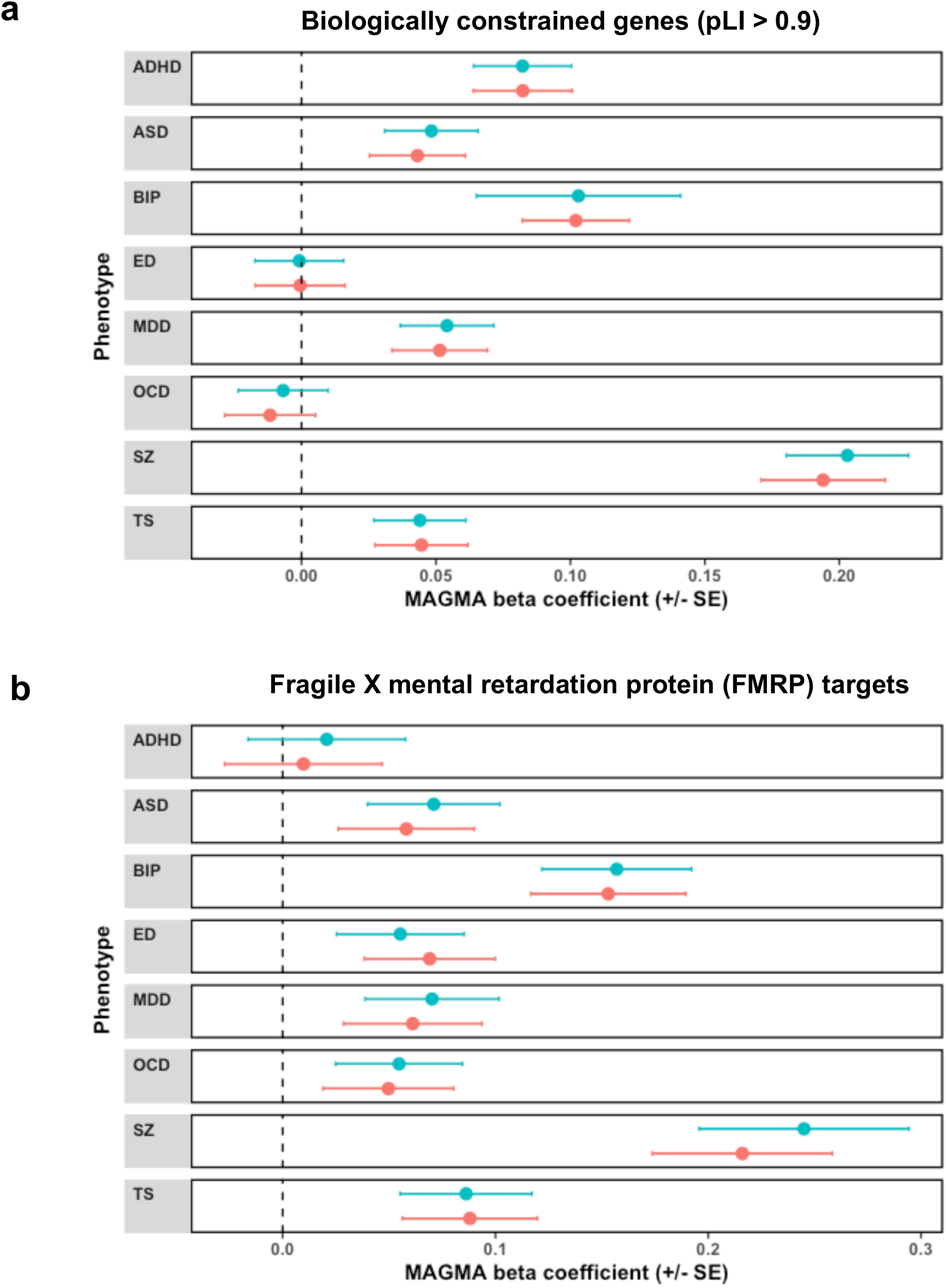
Enrichment of common variant associations within biologically constrained genes and FMRP target genes across different psychiatric disorders. Beta coefficient from MAGMA gene-set association model presented for each psychiatric phenotype, which tested association with (**a**) a set of genes which display intolerance to loss of function variation (probability of loss of function intolerant [pLI] ≥ 0.9, N_Genes_ = 3230), (**b**) targets of FMRP, N_Genes_ = 842. Error bars represent the upper and lower bounds of each coefficient relative to its standard error. Blue points indicative of the MAGMA model constructed without covariation for the expression of each gene in the cortex, whilst pink represents the models covaried for cortical gene expression.

Schizophrenia displayed the strongest association with biologically constrained genes (*β* = 0.203, *SE* = 0.0227, *P* = 1.94 × 10^−19^), followed by BIP (*β* = 0.103, *SE* = 0.0196, *P* = 7.24 × 10^−9^), and ADHD (*β* = 0.082, *SE* = 0.0182, *P* = 3.15 × 10^−6^). Genic constraint has previously been demonstrated to be related to gene expression – thus, as psychiatric risk genes are likely to have high neurological expression we repeated these analyses adjusted for gene-wise expression in the brain (1, 20). After covariation for the expression of each gene in the cortex, this signal remained at least nominally significant in all disorders except ED and OCD (Supplementary Table 7). BIP and schizophrenia demonstrated a notably strong signal for enrichment within the FMRP targets (Schizophrenia: *β* = 0.245, *SE* = 0.0352, *P* = 1.15 × 10^−9^, BIP: *β* = 0.157, *SE* = 0.0493, *P* = 3.93 × 10^−6^). The remaining phenotypes all demonstrated nominal association with the exception of ADHD, with only a minor effect of adjustment for gene-wise brain expression analogous to the models for the biologically constrained gene-set (Supplementary Table 8).

Gene-set association analysis was then undertaken using 7296 hallmark, canonical, and gene ontology (GO) gene-sets from MSigDB. Fifteen of these gene-sets were associated with schizophrenia after Bonferroni correction (*P* < 6.8 × 10^−6^, Supplementary Table 9). The three most significant gene-sets were as follows: *Neuron projection* (*β* = 0.223, *SE* = 0.0383, *P* = 3.23 × 10^−9^), *Somatodendritic compartment* (*β* = 0.254, *SE* = 0.0452, *P* = 9.58 × 10^−9^), and *High voltage gated calcium channel activity* (*β* = 2.1, *SE* = 0.376, *P* = 1.09 × 10^−8^). When gene-set association was performed for the remaining seven phenotypes, none of the fifteen schizophrenia associated gene-sets survived Bonferroni correction for any disorder (Supplementary Table 10).

### Pairwise meta-analysis of schizophrenia with additional psychiatric disorders revealed novel gene-level associations

As there is evidence of wide spread pleiotropy amongst genic associations with schizophrenia in relation to other psychiatric phenotypes, we aimed to further characterise shared common variant genomics using meta-analysis weighted by the sample size of individual GWAS. Schizophrenia was meta-analysed at gene-level in a pairwise fashion with the remaining psychiatric GWAS in order to identify genes which, i) survived multiple testing correction (*P*_Meta_ < 2.7 × 10^−6^) in each meta-analysis but did not survive correction in the respective individual GWAS (*P* < 0.05, *P* > 2.7 × 10^−6^). All seven meta-analyses of schizophrenia and one other psychiatric disorder revealed at least one sub-threshold pleiotropic gene which satisfied the above criteria (Supplementary Table 11, Supplementary Figure 1). The schizophrenia/BIP model yielded the largest number of novel genic associations (N_Novel_ = 175, Lowest *P*: *TSNAXIP1, P* = 2.83 × 10^−10^ – Fig. 3a). Thereafter, the schizophrenia/ADHD (N_Novel_ = 65, Lowest *P*: *ARTN, P* = 1.6 × 10^−9^ – Fig. 3b) and schizophrenia/ASD (N_Novel_ = 58, Lowest *P*: *CXXC4, P* = 2.04 × 10^−8^ – Fig. 3c) meta-analyses had the most novel genic associations. Several previously postulated psychiatric risk genes from candidate studies were revealed in these constructs, including Neural cell adhesion molecule 1 (*NCAM1*) which was significant in the schizophrenia and BIP model (22, 23) and the Delta opioid receptor gene *OPRD1* which survived correction in the schizophrenia meta-analyses with ADHD and MDD respectively (24, 25).

**Figure 3.**
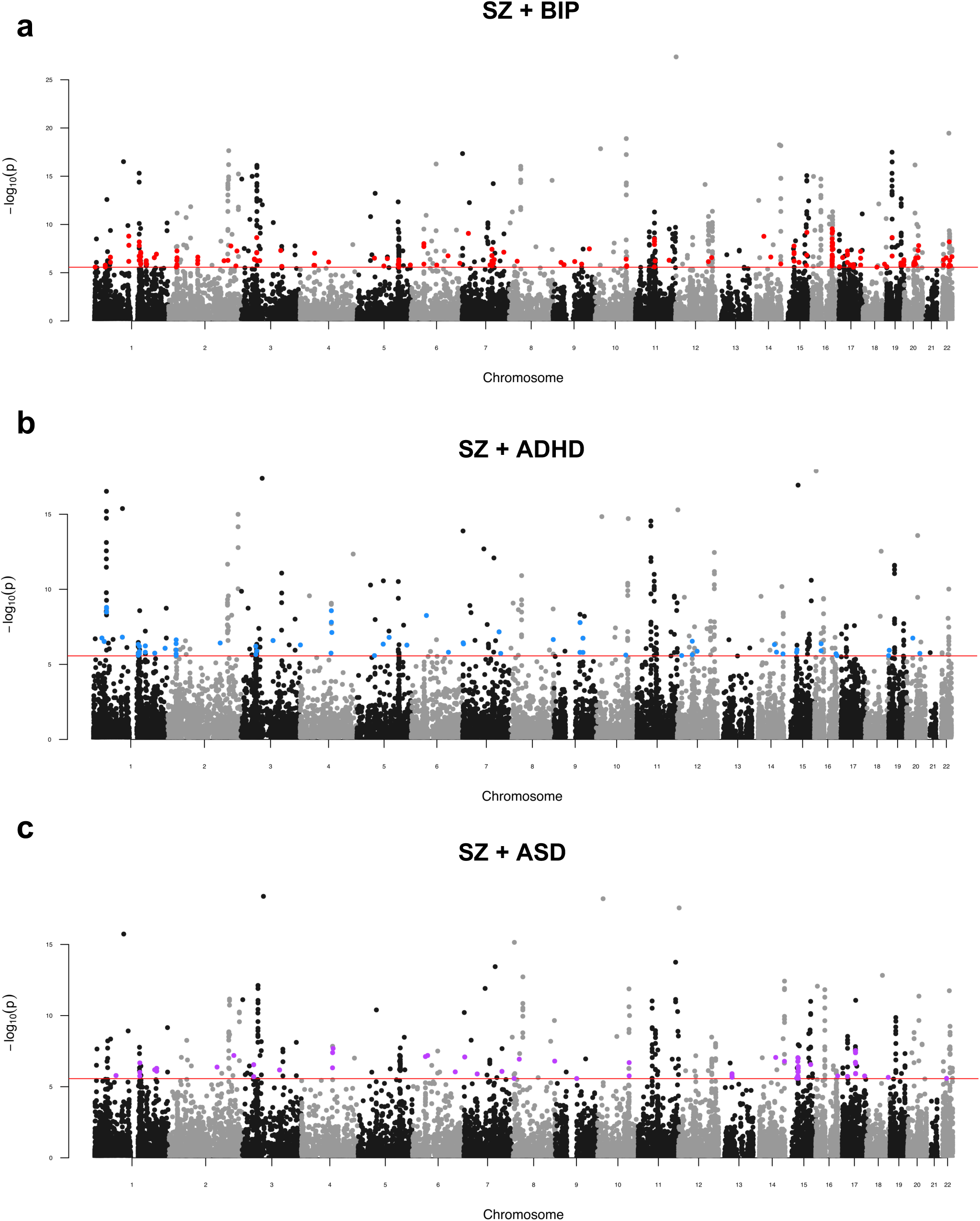
Pairwise genic meta-analysis of schizophrenia and other psychiatric disorders. Manhattan plot for each meta-analysis which displays the −log_10_ transformed *P* value for association for genes which were tagged by at least one SNP in the respective GWAS. The red line represents the Bonferroni threshold for multiple testing correction (*P* < 2.7 × 10^−6^). Genes highlighted on each plot were not Bonferroni significant in the individual GWAS but obtained corrected significance in the meta-analysis. (**a**) Schizophrenia (SZ) and Bipolar Disorder (BIP) genic meta-analysis, (**b**) Schizophrenia and Attention Deficit/Hyperactivity Disorder (ADHD) genic meta-analysis. (**c**) Schizophrenia and Autism Spectrum Disorder (ASD) genic meta-analysis. Manhattan plots for the meta-analyses with MDD, TS, ED, and OCD are presented in supplementary figure 1.

Across all seven meta-analyses there were 330 genes in total which only achieved corrected significance once schizophrenia was combined with another psychiatric trait. A notable proportion of this subset (N = 76) were identified across multiple meta-analyses, of which 20 survived correction in three pairwise models – for example, *CXXC4* which is a regulator of Wnt signalling (26). Gene-set association analysis used the *Z*-scores derived from each meta-analysis for the data-driven gene-sets from MSigDB. All pairwise models demonstrated at least one gene-set which survived multiple testing correction, largely recapitulating the gene-sets which were also significant in the univariate analysis of schizophrenia (Supplementary Table 12).

## DISCUSSION

We investigated the common variant genomic architecture of schizophrenia relative to a range of psychiatric disorders. Previous work has suggested that there is a significant relationship between the genetics of schizophrenia and the spectrum of psychiatric phenotypes (27, 28), which is not unexpected given the analogous aspects of their respective clinical presentations. Gene-based analysis consolidated this aetiological overlap as schizophrenia shared several significant genic associations with other psychiatric disorders. MDD and BIP demonstrated the largest number of shared schizophrenia genes relative to the total number of genes significant for each disorder, consistent with the strong genetic correlation between schizophrenia and these phenotypes. Interestingly, the most strongly associated schizophrenia gene in this study, *CACNA1C*, was also strongly enriched with common variant associations for BIP. The Cav1.2 L-type voltage-gated calcium channel α1c subunit is encoded by *CACNA1C*, which is postulated to play an important role in synaptic plasticity (29-31).

The majority of schizophrenia associated genes were not significant for any other disorder after the application of multiple testing correction. However, a large proportion of these genes displayed some degree of nominal significance (*P* < 0.05) – thus, we identified a subset of schizophrenia genes which displayed no evidence of association with any other psychiatric disorder (*P* > 0.05). This strict threshold was chosen as the sample sizes differ amongst the respective GWAS and future increases in sample sizes may yet reveal nominal *P* values which survive multiple testing correction. The biological saliency of variation mapped to a subset of these genes was supported by TWAS. For instance, decreased predicted expression of the potassium channel *KCNN3* was associated with schizophrenia. Dopamine homeostasis has been extensively linked to this protein, with blockade shown to increase spike firing of dopaminergic neurons, along with potentiated dopamine release (32, 33). Downregulation of *KCNN3*, therefore, is consistent with overarching dopamine hypotheses related to schizophrenia, supporting its relevance for pathogenesis of the disorder.

The enrichment of common variant associations within genes under biological constraint was shown to be relatively ubiquitous across most of the psychiatric phenotypes considered in this study. These mutation intolerant regions of the genome are subject to purifying selection and, thus, it may appear contradictory that risk alleles for psychiatric disorders persist in these genes at common frequencies. It has been postulated that linkage between regions under repeated selection results in the loss of haplotypes, in turn, attenuating the strength of selection on individual sites (1, 34, 35). This weakened selective pressure is theorised to account for the increase in frequency of these risk alleles through the action of enhanced genetic drift (34, 36). Therefore, this mechanism of common variant enrichment within constrained regions appears to be an important aspect of the genomic architecture of several psychiatric disorders which is shared with schizophrenia. OCD did not display a significant association with this gene-set, as perhaps would be expected, however, the small sample size of its GWAS may account for this. Deleterious rare variation within these regions has also shown to be enriched for schizophrenia and ASD, consolidating the multi-faceted nature of genomic risk for these disorders (3, 37).

This study refines the nature of common variant informed genetic overlap between schizophrenia and other psychiatric conditions. A number of interesting pleiotropic candidates were revealed in this study which warrant further functional investigation to elucidate their significance to the respective phenotypes. For instance. *SORCS3*, the most pleiotropic schizophrenia-associated gene uncovered by these analyses due to its association with two of the other disorders considered, is implicated in a number of neurologically salient processes including modulation of synaptic depression and glutamate receptor functionality (38, 39). There are a number of important limitations to this study. Firstly, this work relies on the diagnostic definitions encompassed within each GWAS, however, the true incidence of psychiatric comorbidities amongst the respective study participants is unknown. Despite the uncertainty of psychiatric nosology, schizophrenia associated genes involved with another psychiatric GWAS does suggest that these shared genes may be particularly biologically salient. Further, the non-pleiotropic subset of genes identified in this study may be associated with other psychiatric phenotypes not considered in this study. We utilised common variant data for this study. Whilst common variation is an integral component of psychiatric heritability, future cross-disorder investigation of genes and systems significantly enriched with rare loci will be integral to fully appreciate the spectrum of biology which constitutes the shared factors which exist between schizophrenia and these disorders. Genic coordinates for SNP annotation was also largely constrained to protein coding genes, however, previous evidence has suggested that non-coding RNA play an important role in psychiatric disorders, with variation mapped to the microRNA host gene *MIR137HG* a notable significant GWAS signal for schizophrenia (40). Future study should consider if there are shared association signals mapped to these non-coding RNA, particularly as transcriptomic imputation has not yet thoroughly applied to small RNA sequencing data. European GWAS data were chosen for these analyses as diverse well-powered summary statistics are still not readily available for many psychiatric traits. There are important ancestral differences between haplotype structure, and thus, it is essential that GWAS are more widely performed for non-European ancestral groups so that work of this fashion can be consolidated and leveraged in an inclusive fashion. Finally, the principal multimarker test employed for gene discovery in this study, MAGMA, is based on *P*-value combination and assigns variants to genes using their genomic coordinates. This does not directly inform the effect size or functional significance of variation which constitutes these gene-based *P* values. TWAS helps to overcome this by assigning weights to SNPs based on the *cis*-heritability of their respective genes. However, TWAS relies on genotyped expression datasets of modest sample sizes, with many genes lacking a suitable imputed model of genetically regulated expression. Further study is required to advance these multimarker approaches such that statistical effect size and functional annotation beyond *cis*-regulatory annotations can be included. Enhanced models which integrate this information will facilitate the biological interpretation of schizophrenia associated genes shared with other disorders

## MATERIALS AND METHODS

### GWAS summary statistics

The largest European ancestry GWAS with full genome-wide summary statistics available was obtained for schizophrenia (N = 105318, Pardiñas *et al.* (1)) and seven other psychiatric disorders. The other disorders and their respective GWAS were as follows: bipolar disorder (BIP, N = 51710) (8), attention-deficit/hyperactivity disorder (ADHD, N = 53293 [European subset]) (9), major depressive disorder (MDD, N=173005) [excluding the 23andMe cohorts for which full summary statistics are not publicly available]) (10), obsessive compulsive disorder (OCD, N = 9725) (11), eating disorder (ED, N=14477) (12), autism spectrum disorder (ASD, N=46351) (13), and Tourette’s Syndrome (TS, N=14307) (14). Schizophrenia summary statistics were downloaded from the Walters group data repository (https://walters.psycm.cf.ac.uk/), whilst the other seven summary statistics were obtained from the website of the psychiatric genomics consortium (PGC, https://www.med.unc.edu/pgc/results-and-downloads/).

### Gene-based association analysis

Gene-based association was undertaken for each of the disorders using the MAGMA package v1.06b (mac) as described elsewhere (41). Briefly, the gene-based method implemented by MAGMA utilises *P*-values as input, whereby the test statistic is a linear combination of genic *P*-values. We used the default gene-based test in MAGMA which was the mean of the *χ*^2^ for the variants annotated to each gene. In order to account for dependent *P*-values due to linkage disequilibrium (LD) between variants, the 1000 genomes phase 3 European reference panel is used to derive variant-wise LD such that the null distribution can be approximated. Variants were mapped to 18297 autosomal protein-coding genes in hg19 genome-assembly (NCBI), obtained from the MAGMA website (https://ctg.cncr.nl/software/magma). We removed genes which arise from the extended major histocompatibility complex (MHC) on chromosome 6 due to the complexity of LD in that region, as is standard practice. Genic coordinates were extended 5 kilobases (kb) upstream and 1.5 kb downstream during annotation to capture potential regulatory variation. Statistical inference for a significantly associated gene for each disorder was set as *P* < 2.7 × 10^−6^ to adjust for the number of genes tested via the Bonferroni method. Genes associated with schizophrenia which did not display nominal association with any of the other disorders (*P* < 0.05) were subject to overrepresentation analysis (ORA) using the entire set Bonferroni significant schizophrenia genes as the statistical background. ORA was performed by ConsensusPathDB (http://cpdb.molgen.mpg.de/) (42). A minimum overlap of three input genes was selected, with analysis utilising pathway-based sets and level two gene-ontology sets.

### Gene-set association analysis

Competitive gene-set association for each disorder was undertaken with MAGMA. This model tests the underlying null hypothesis that genes within a set are no more strongly associated with the phenotype than all other genes (41, 43). MAGMA constructs a linear regression model wherein genic-association (transformed to *Z* via the probit function) is the outcome, with adjustment for confounders including gene-size and genic-minor allele count. A one-sided test is performed for the term in the model which specifies whether each gene is within the set of interest (*β*_*GS*_), such that the null hypothesis is *β*_*GS*_ = 0 and the alternative *β*_*GS*_ > 0. We selected 7296 hallmark, canonical, and gene ontology genesets from the molecular signatures database (MSigDB) for gene-set association (19). We also tested two hypothesis driven gene-sets which survived multiple testing correction in the Pardiñas *et al.* schizophrenia GWAS analyses (1). Probability of loss of function intolerance (pLI), as a metric of biological constraint, was obtained from ExAC – with all genes displaying high intolerance (pLI ≥ 0.9, N_Genes_ = 3230) were included in the first set (20). As constrained genes are postulated to have increased expression in the brain (20), we also repeated the analysis covaried for the expression of each gene (median transcript per million) in the brain. Genes were annotated using RNA-sequencing data for the brain (cortex) from the genotype-tissue expression consortium (GTEx v7) (44). The second hypothesis driven set was obtained from Darnell *et al.*, selecting FMRP targets using a stringent false discovery rate (FDR) < 0.01 threshold (21).

### Transcriptome-wide association studies (TWAS)

TWAS was performed using FUSION (https://github.com/gusevlab/fusion_twas/) with default settings as described previously (17, 18). Summary statistics were formatted using the LDSC framework as is usual practice (munge_sumstats.py, https://github.com/bulik/ldsc) (45). SNP weights were obtained from the FUSION website for models imputed using data from the DLPFC (CommonMind Consortium) and blood (Young Finns Study) (46, 47). Statistical inference, to account for the number of models tested, was set at *P* < 9.22 × 10^−6^ (5420 models tested) and *P* < 1.06 × 10^−5^ (4701 models tested) for the DLPFC and blood respectively. The underlying prinicipal of TWAS is that the expression of genes identified through this method are genetically correlated with the phenotype of interest. Conditional analysis was then undertaken to identify disorder associated genes which are independent versus those co-expressed with a genetic predictor which is shared between them. This was implemented with the FUSION package R script FUSION.post.process with the locus window set at 100000 base pairs.

### Pairwise cross-disorder meta-analysis

Genic *Z* scores for schizophrenia were meta-analysed in a pairwise manner with the other seven disorders using the --meta flag in MAGMA. This utilises Stouffer’s weighted *Z* method, in which the *i*^*th*^ *Z* score is weighted (*w*_*i*_) by the sample size of the respective GWAS:

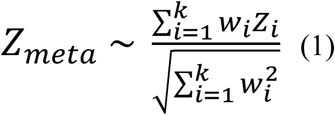

Analogous to the univariable gene-based method, the test-statistic follows a standard normal distribution under the null hypothesis (48). Gene-set association was then undertaken using the model described above with *Z*_*meta*_ the response variable in each instance.

## Code and data availability

Summary statistics are publicly available as described in the GWAS summary statistics section of the materials and methods. All software is publicly available with versions and specific scripts also described in the materials and methods. Command line inputs and code for figure generation in this study are also available (
https://github.com/Williamreay/Schizophrenia_cross_disorder_scripts)

