## Supplementary Figure for "Pairwise common variant meta-analyses of schizophrenia with other psychiatric disorders reveals shared and distinct gene and gene-set associations"

**a**

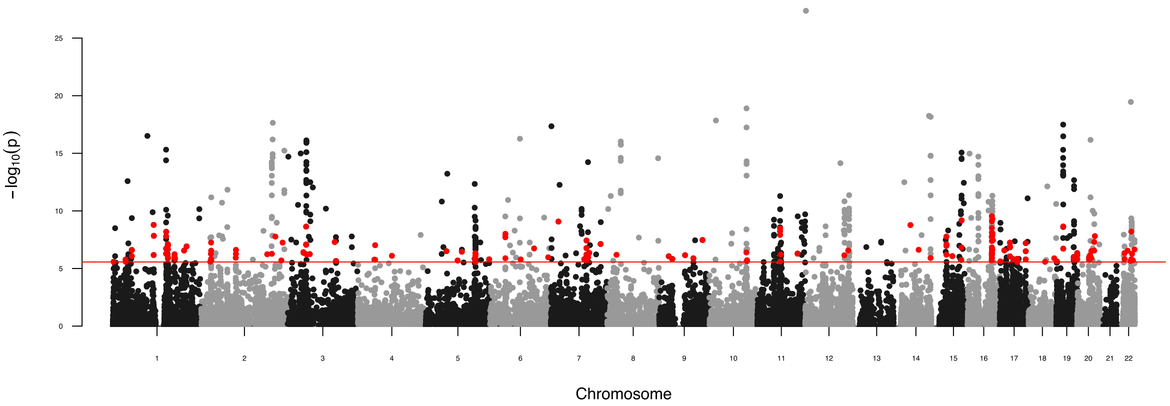

**b**

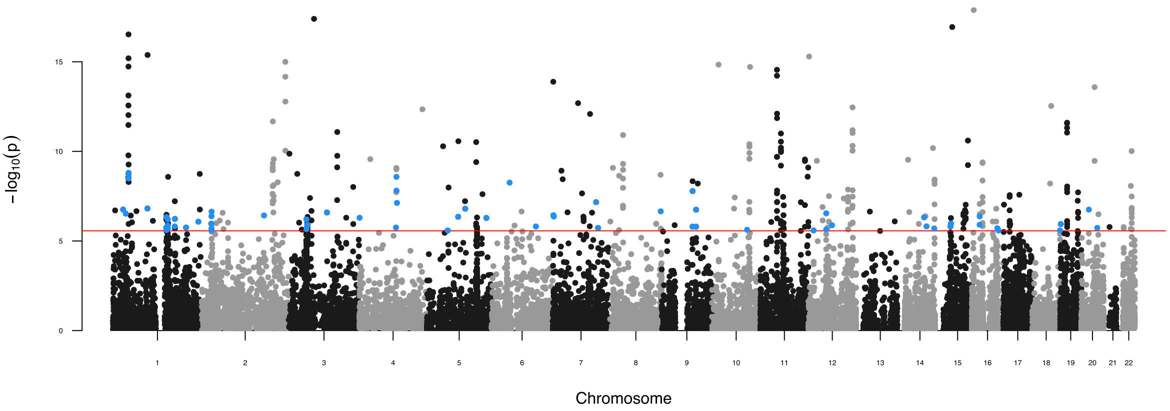

**c**

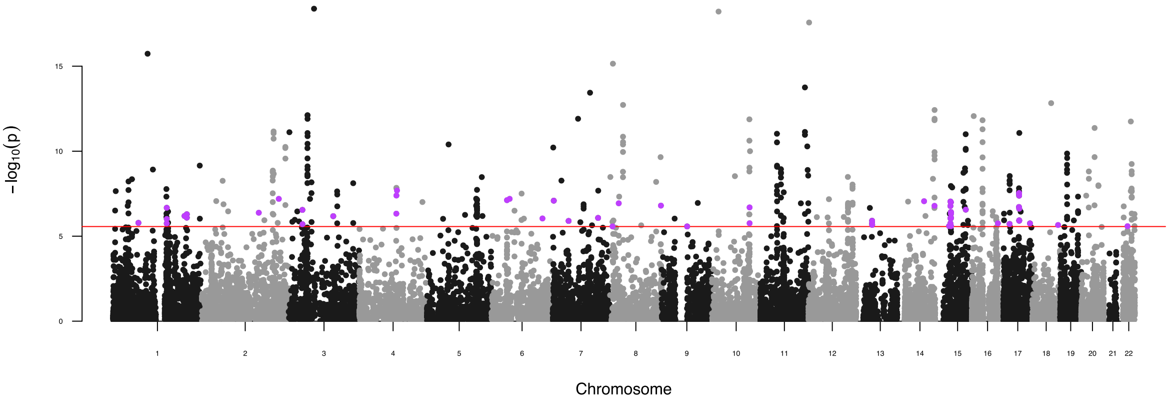
−log10(p)
0
5
10
15
Chromosome

**d**

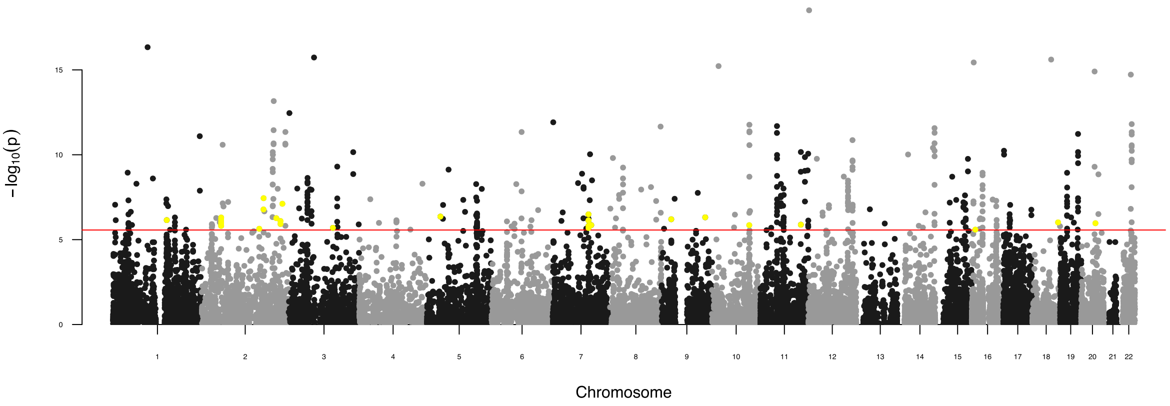
−log10(p)
0
5
10
15
Chromosome

**e**

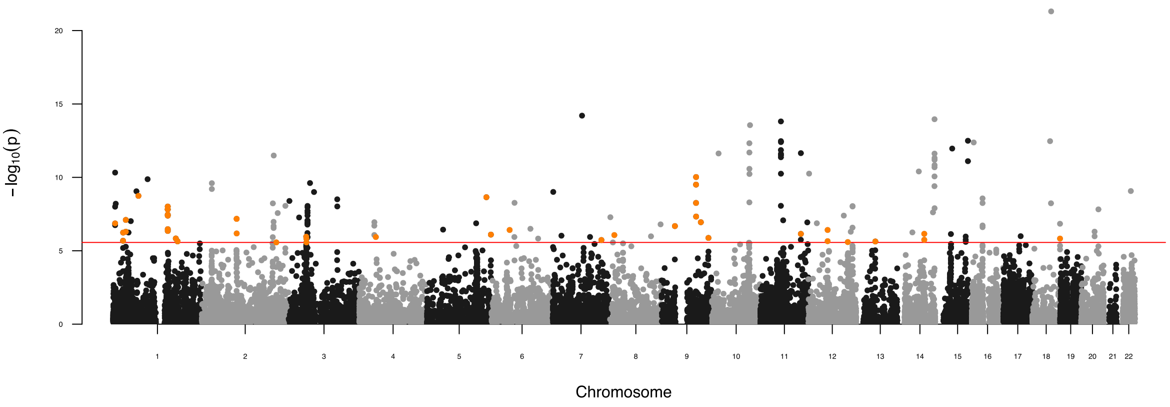
−log10(p)
0
5
10
15
20
Chromosome

**f**

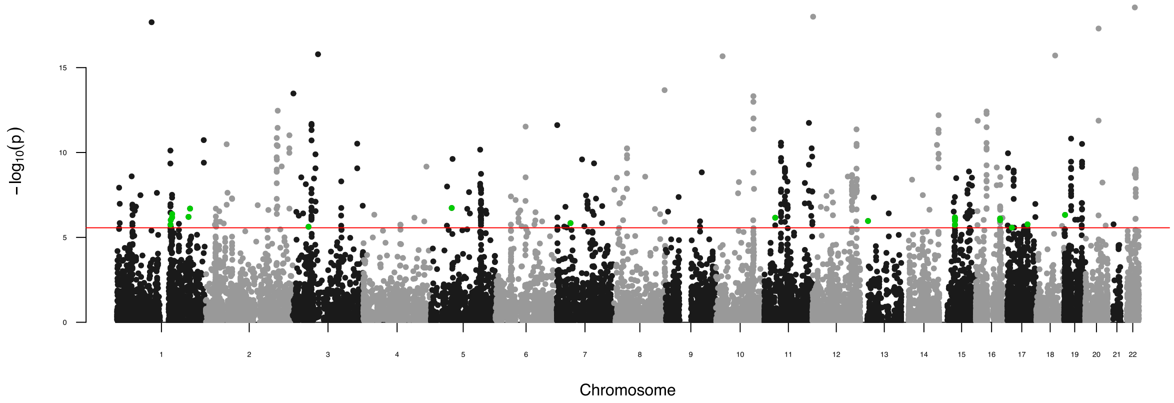

**g**

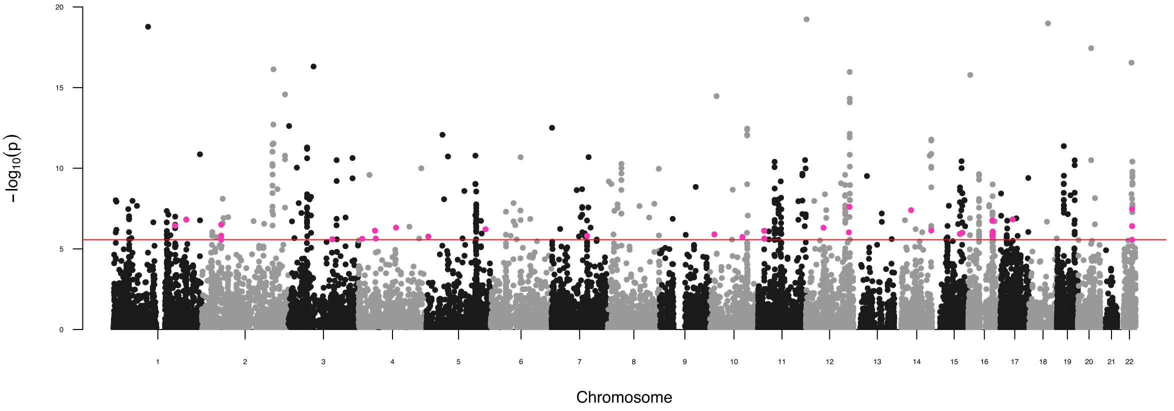

**Supplementary Figure 1. Pairwise genic meta-analysis of schizophrenia and other psychiatric disorders.** Manhattan plot for each meta-analysis which displays the -$\log_{10}$ transformed *P* value for association for genes which were tagged by at least one SNP in the respective GWAS. The red line represents the Bonferroni threshold for multiple testing correction ($P < 2.7 \times 10^{-6}$). Genes highlighted on each plot were not Bonferroni significant in the individual GWAS but obtained corrected significance in the meta-analysis. (**a**) Schizophrenia (SZ) and Bipolar Disorder (BIP) genic meta-analysis, (**b**) Schizophrenia and

Attention Deficit/Hyperactivity Disorder (ADHD) genic meta-analysis. (**c**) Schizophrenia and Autism Spectrum Disorder (ASD) genic meta-analysis, (**d**) Schizophrenia and Eating Disorder (ED) genic meta-analysis, (**e**) Schizophrenia and Major Depressive Disorder (MDD) genic meta-analysis, (**f**) Schizophrenia and Obsessive-Compulsive Disorder (OCD) genic meta-analysis, (**g**) Schizophrenia and Tourette's Syndrome (TS) meta-analysis.
